# Region-Specific Changes in the Aging Enteric Nervous System

**DOI:** 10.64898/2026.11.17.767400

**Authors:** Jack H. Marciano, Shaul Druckmann, Laren S. Becker, Julia A. Kaltschmidt

## Abstract

As human life expectancy increases, understanding the physiological changes that accompany aging becomes increasingly important. Gastrointestinal (GI) dysfunction is common with age, and age-related changes in the enteric nervous system (ENS), which regulates GI motility, may contribute to this decline in function. Studies of the aging ENS have reported contradicting results, potentially reflecting the substantial regional heterogeneity of the ENS and the fact that most studies focus on only one or two areas of the GI tract. Neurons in the myenteric plexus, which regulate intestinal contractions and motility, are organized into ganglia that form circumferential stripes along the length of the gut. Here, we systematically compare neuronal and glial density and organization across all five regions of the small and large intestine in young (3-4 month-old) and aged (22-26 month-old) mice. We find that neuronal density remains unchanged with age across all regions, despite localized changes in ganglia organization. Although the overall stripe-like arrangement is preserved, ganglia are less aligned with stripes in the duodenum and ileum, while those in the ileum and distal colon are smaller, contain fewer neurons, and have a greater proportion of neurons located outside the ganglia. Glial cell numbers vary regionally with age, increasing in the duodenum and distal colon but decreasing in the proximal colon. The proportion of Calretinin^+^ neurons, meanwhile, remains stable across the small intestine. Together, these findings reveal that aging is associated with selective, region-specific remodeling of myenteric plexus cytoarchitecture rather than uniform structural decline across the GI tract. This comprehensive regional analysis may help reconcile conflicting reports of age-related changes in the ENS.

## 1 Introduction

Age-associated diseases and physiological dysfunctions represent an increasingly pressing burden on human health. Gastrointestinal (GI) dysfunction is common with aging and is associated with changes in GI motility and an increased prevalence of GI disorders (Dumic et al., 2019). The enteric nervous system (ENS) is a semi-autonomous network of neurons and glia embedded within the GI tract and regulates many aspects of GI motility, secretion, and blood flow (Furness, 2012; Avetisyan et al., 2015). Given its central role in coordinating these functions, the ENS has been implicated as a potential contributor to age-related GI dysfunction, with age-associated changes in enteric neuronal number, composition, and organization reported in both human and rodent studies (Nguyen et al., 2023). However, a region-wide map of how the ENS ages across the different gut regions has not yet been established.

The GI tract is a highly heterogeneous organ, with distinct regions specialized for different physiological functions (Hornbuckle et al., 2008). The ENS similarly exhibits substantial regional variation in its cytoarchitecture and cellular composition. It is organized into two major plexuses: the myenteric plexus is positioned between the longitudinal and circular muscle layers and primarily regulates GI motility, while the submucosal plexus regulates epithelial secretion, absorption, and blood flow (Furness, 2012). Within the myenteric plexus, enteric neurons and glia are organized into distinct ganglia interconnected by nerve fibers, with ganglion density and organization varying across the length of the GI tract (Sasselli et al., 2012). At a larger spatial scale, myenteric ganglia form a stripe-like pattern, with both stripe width and inter-stripe distance varying between gut regions (Hamnett et al., 2022). Neuronal subtype composition also varies substantially across the GI tract (Hamnett et al., 2022). Thus, the ENS is not a homogeneous neural network, but instead exhibits pronounced regional differences in its cellular composition and spatial organization. This anatomical heterogeneity raises the possibility that age-associated changes in ENS structure may differ across the GI tract.

Most studies on ENS aging have focused on analyzing neuronal number and density in the myenteric plexus of relatively restricted areas of the GI tract. While some studies in mice and rats find neuronal loss (Santer, 1994; El-Salhy et al., 1999; Phillips et al., 2004), others find no significant change in neuronal number (Gamage et al., 2013). Reports further describe reduced packing density of aged myenteric ganglia, swollen, dystrophic axonal profiles, and a particular vulnerability of neurons expressing calcium-binding proteins (El-Salhy et al., 1999; Gamage et al., 2013; Wang et al., 2013; Becker et al., 2018). One study that has examined multiple GI regions in aged rats suggests greater neuronal loss in the colon than in the small intestine (Phillips et al., 2004). However, these studies have largely focused on neuronal number and do not examine other features of ENS cytoarchitecture, including neuronal stripe architecture, ganglionic patterning, and glial organization. Thus, the extent and regional specificity of age-associated ENS remodeling across the GI tract remain unclear. Here, we performed a systematic analysis of myenteric neuronal cytoarchitecture across five anatomically distinct regions of the small and large intestine in young (3-4 month-old) and aged (22-26 month-old) mice. Using immunohistochemistry and quantitative image analysis, we examined neuronal stripe architecture and ganglionic organization. We also assessed glial number and organization, as well as the proportion of Calretinin^+^ neurons, a population that includes excitatory motor neurons.

We found that neuronal density was unchanged with age, but ENS cytoarchitecture exhibited region-specific changes. Ganglia were less aligned with the stripe pattern in the duodenum and ileum and ganglionic size was altered in the ileum and distal colon. Glial cell numbers also changed with age, increasing in the duodenum and distal colon and decreasing in the proximal colon. These findings demonstrate that aging does not uniformly alter ENS architecture throughout the GI tract but instead produces spatially heterogeneous remodeling.

## 2 Results

To determine whether the organization of the myenteric plexus changes with age, and whether such changes differ between gut regions, we compared young (3-4 month-old) and aged (22-26 month-old) male mice across five regions of the small and large intestine: the duodenum, jejunum, ileum, proximal colon, and distal colon (Supplementary Figure S1A). We focused on the myenteric plexus because it contains the intrinsic neuronal circuitry required for GI motility and its organization has been extensively characterized in young mice (Spencer and Hu, 2020; Hamnett et al., 2022). To label enteric neurons, we used the pan-neuronal marker HuC/D.

We first asked whether aging alters the large-scale spatial organization of the myenteric plexus neurons. In young mice, myenteric ganglia are arranged in a stripe-like pattern, with stripe width and spacing varying between gut regions (Hamnett et al., 2022). The number of HuC/D^+^ neurons per field did not differ between young and aged mice in any region (Figure 1C). To determine whether these neurons change their spatial arrangement, we identified HuC/D^+^ neurons using a modified version of COUNTEN (Kobayashi et al., 2021) and grouped them into ganglia, defined as clusters of 3 or more neurons (see Methods). We then traced the stripes in each field blind to age, and measured how closely the long axis of each ganglion followed the stripe direction, using ganglia of ≥8 neurons, whose orientation can be reliably measured (Figures 1A,B; see Methods). Stripes could be identified in every field at both ages, and in young mice the long axis of their ganglia was well aligned with stripes in all regions (mean alignments of 0.46–0.66, on a scale from –1, perpendicular, to +1, parallel). With age, alignment significantly decreased in the ileum, from 0.53 to 0.19, and to a lesser extent in the duodenum, from 0.66 to 0.42, but did not change in the jejunum, proximal colon, or distal colon (Figure 1D). Thus, the overall stripe-like organization of the myenteric plexus is preserved with age, although ganglia become less aligned with this pattern in the duodenum and ileum.

**Figure 1.**
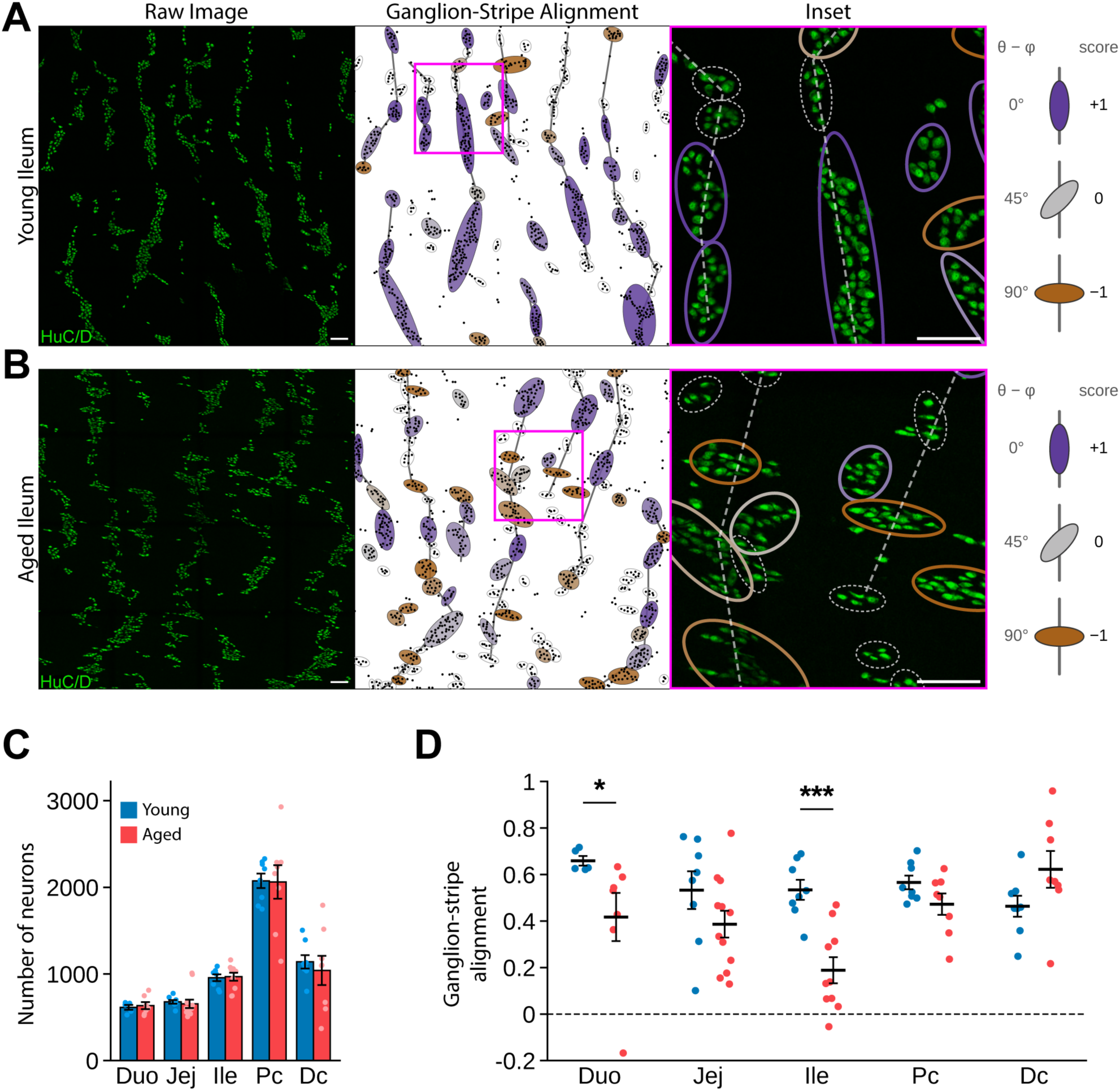
Ganglia become less aligned to the stripe pattern of the myenteric plexus with age in a region-specific manner. **(A, B)** Representative HuC/D^+^ neuronal immunostaining (green, left column) in the ileum of a young **(A)** and aged **(B)** mouse, with the corresponding ganglion-stripe alignment maps (middle column) and enlargements of boxed regions (right column). In the alignment maps, neurons are shown as black dots and stripes, traced blind to age, as grey lines. Each ganglion of ≥8 neurons is shown as an ellipse fitted to its outline and colored by its alignment with the stripe direction of the field, from parallel (+1, purple) to perpendicular (–1, brown); ganglia of 3-7 neurons were outlined in grey, and were not scored (see Methods). In the enlargements, ellipses are overlaid on the HuC/D^+^ immunostaining. **(C, D)** Quantification of the number of neurons per 3.24 mm^2^ field **(C)** and of ganglion-stripe alignment **(D)** across all gut regions. Data are presented as mean ± SEM, with individual animals shown as points; *n* = 5-12 mice per group (8 young and 12 aged in total). (C, D) Linear mixed models followed by within-region comparisons of age; D was weighted by the number of scored ganglia per mouse and region. *p<0.05, **p<0.01, ***p<0.001. Scale bars, 100 µm. Duo, Duodenum; Jej, Jejunum; Ile, Ileum; Pc, Proximal colon; Dc, Distal colon.

We next examined ganglion-level organization to determine whether aging also alters how neurons are distributed between ganglia and the surrounding plexus. Using the same ganglion definition of 3 or more neurons, we classified neurons outside these clusters as extraganglionic (Figures 2A,B; Supplementary Figure S1B; see Methods). The total number of ganglia did not change in any region when comparing young and aged mice (Figure 2C). However, in the ileum and distal colon of aged mice, the number of extraganglionic neurons increased by 50% and 47%, respectively, and the percentage of neurons that were extraganglionic rose from 13.0% to 19.3% in the ileum and from 11.7% to 23.4% in the distal colon (Figures 2D,E). In these same regions, mean ganglion area decreased, as did the mean and the 90^th^ percentile of neurons per ganglion (Figures 2F-H). To determine whether a loss in the largest ganglia accounted for this decrease, we binned ganglia by neuron number and compared the largely unaffected duodenum with the ileum (Figures 2I,J). In the ileum, a smaller proportion of ganglia contained ≥10 neurons, whereas the distribution in the duodenum was unchanged. Consistent with this, the ileum and distal colon had a smaller proportion of large ganglia (Supplementary Figures S1C-L). Thus, while total neuron and ganglia numbers are maintained with age, ganglia in the ileum and distal colon are smaller and contain fewer neurons, with a greater proportion of neurons lying outside of ganglia. The duodenum, jejunum, and proximal colon, while trending in a similar direction, are comparatively unaffected.

**Figure 2.**
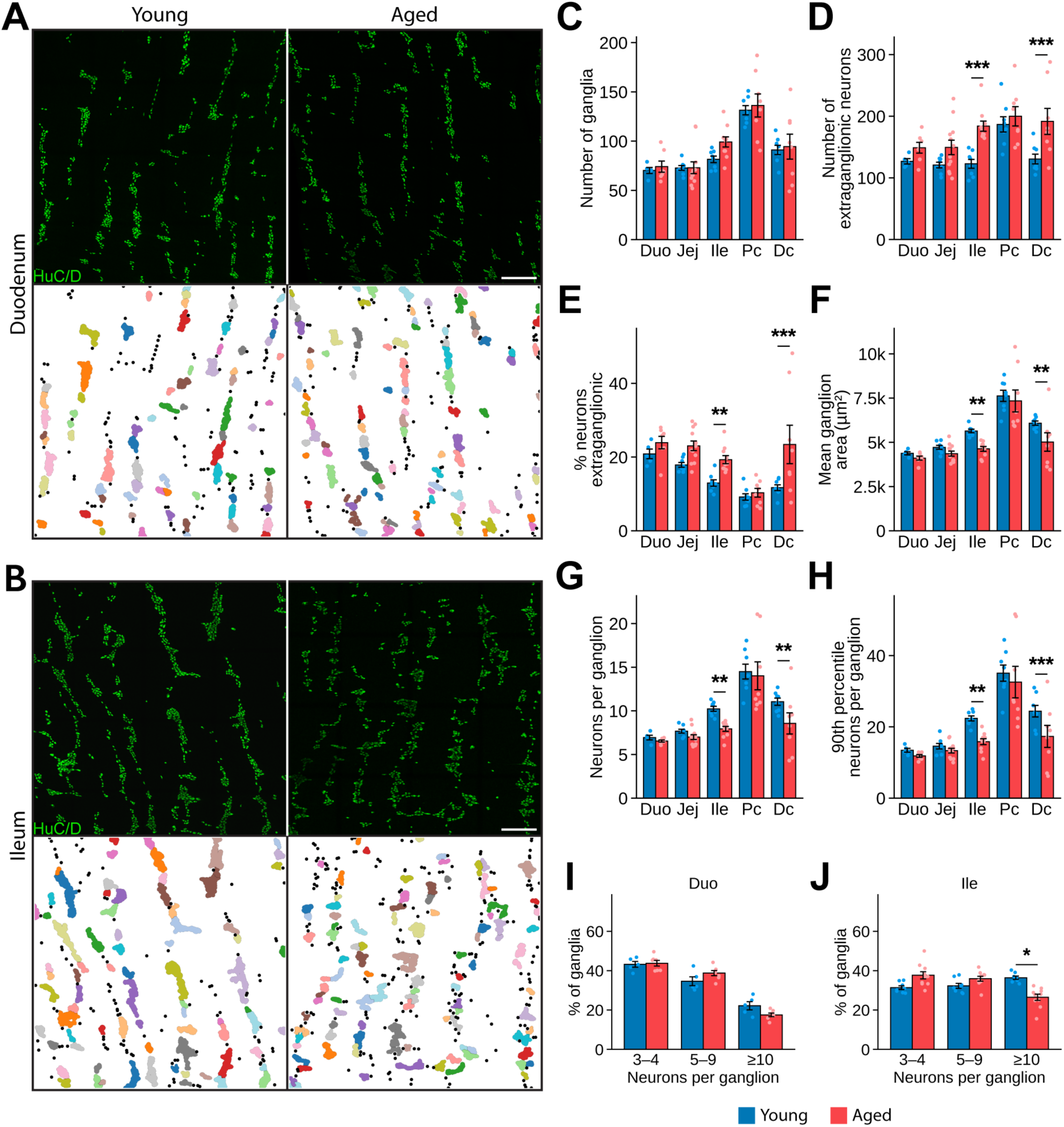
Ganglionic organization shifts with age in a region-specific manner. **(A, B)** Representative HuC/D^+^ neuronal immunostaining (green) in the duodenum **(A)** and ileum **(B)** of young (left) and aged (right) mice, with the corresponding ganglia masks below. Ganglia (clusters of ≥3 neurons, see Methods) are shown in distinct colors and extraganglionic neurons as black dots. **(C-H)** Quantification across all gut regions of the number of ganglia **(C)** and extraganglionic neurons **(D)** per 3.24 mm^2^ field, the percentage of neurons that are extraganglionic **(E)**, and mean ganglion area **(F)**, mean neurons per ganglion **(G)** and 90^th^-percentile neurons per ganglion **(H)**. **(I, J)** Percentage of ganglia containing 3-4, 5-9, or ≥10 neurons in the duodenum **(I)** and ileum **(J)** of young and aged mice. Bins were defined by the tertiles of neurons per ganglion pooled across all animals and regions. Data are presented as mean ± SEM, with individual animals shown as points; *n* = 5-12 mice per group (8 young and 12 aged in total). (C, D, F, G, H) Linear mixed models (F, G, H on log_10_-transformed values; F weighted by number of ganglia) and (E) binomial GLMM, each followed by within-region comparisons of age. *p<0.05, **p<0.01, ***p<0.001. (I, J) Binomial GLMMs followed by within-region comparisons of age with Benjamini-Hochberg correction. *q<0.05. Scale bars, 250 µm. Duo, Duodenum; Jej, Jejunum; Ile, Ileum; Pc, Proximal colon; Dc, Distal colon.

We next asked whether enteric glia were altered with age, and whether any changes follow the same regional pattern as neuronal organization. Enteric glia are found within myenteric ganglia, along the fibers that connect them, and within the smooth muscle layers (Santhosh et al., 2024). To label enteric glia, we used an antibody against Sox10, a transcription factor expressed in enteric glial nuclei (Rao et al., 2015; Figures 3A,B; Supplementary Figures S2A,B). With age, the total number of Sox10^+^ cells increased in the duodenum and distal colon, and decreased in the proximal colon (Figure 3C). To determine whether these changes predominantly occurred within ganglia, we used COUNTEN to generate a ganglion mask from the HuC/D channel of each field and counted the Sox10^+^ cells within it (Supplementary Figure S2C; see Methods). The number of intraganglionic Sox10^+^ cells increased with age only in the distal colon (Figure 3D), while the number of extraganglionic Sox10^+^ cells increased in the duodenum and distal colon and decreased in the proximal colon (Figure 3E). However, the percentage of Sox10^+^ cells that were intraganglionic did not change in any region (Figure 3F). Thus, glial number changes with age in a region-specific manner, increasing in the duodenum and distal colon and decreasing in the proximal colon, while the proportion of glia located within ganglia is maintained.

**Figure 3.**
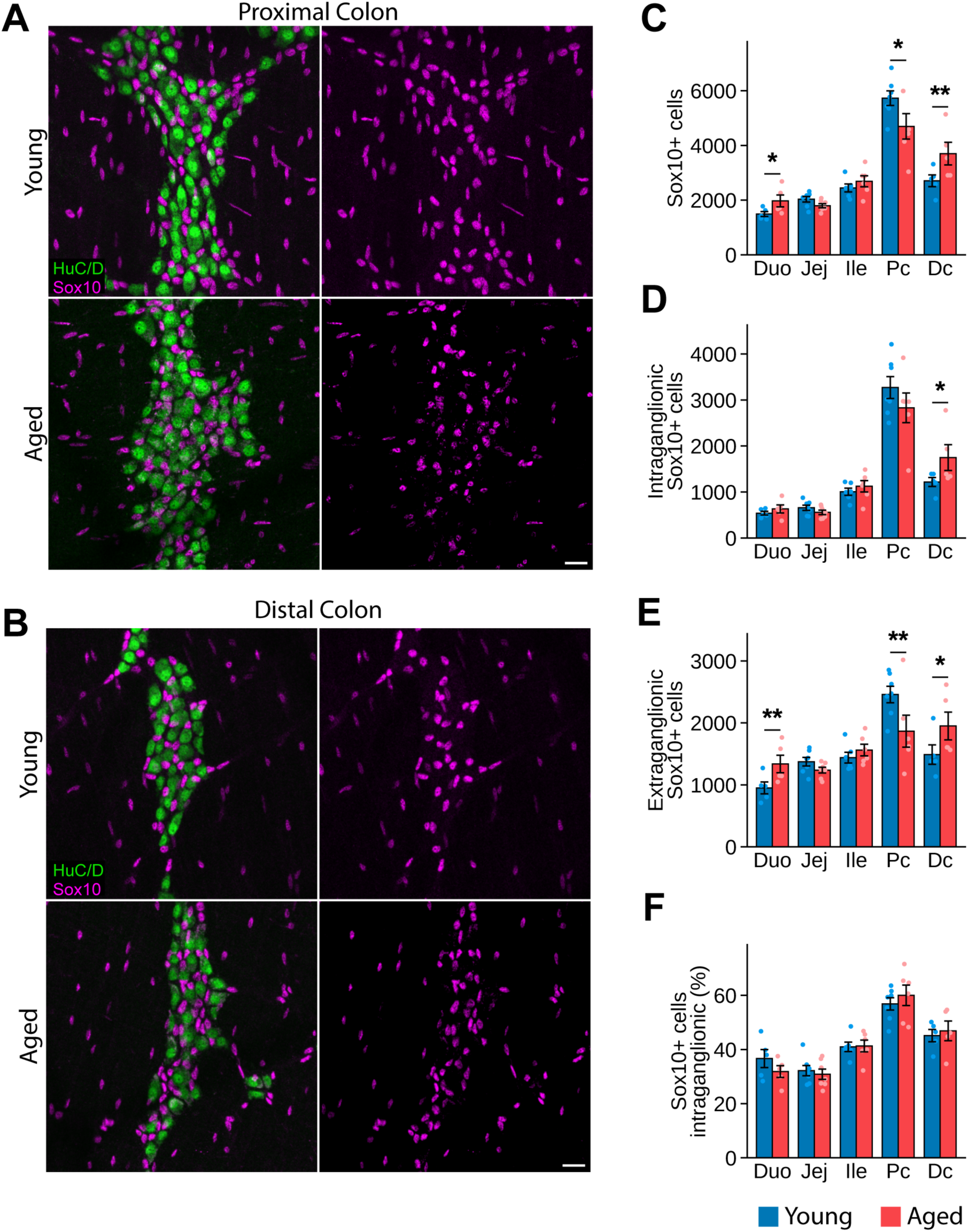
Glial number, but not distribution, changes with age in a region-specific manner. **(A, B)** Representative HuC/D^+^ neuronal (green) and Sox10^+^ glial (magenta) immunostaining in the proximal colon **(A)** and distal colon **(B)** of young (top) and aged (bottom) mice. **(C-F)** Quantification across all gut regions of the number of Sox10^+^ cells **(C)**, intraganglionic Sox10^+^ cells **(D)**, and extraganglionic Sox10^+^ cells **(E)** per 3.24 mm^2^ field, and the percentage of Sox10^+^ cells that are intraganglionic **(F)**. Data are presented as mean ± SEM, with individual animals shown as points, *n* = 5-7 mice per group (8 young and 8 aged in total). Generalized linear mixed models followed by within-region comparisons of age: Poisson for (C-E) and binomial for (F). *p<0.05, **p<0.01, ***p<0.001. Scale bars, 25 µm. Duo, Duodenum; Jej, Jejunum; Ile, Ileum; Pc, Proximal colon; Dc, Distal colon.

Neurons expressing calcium-binding proteins have been reported to be particularly vulnerable to age-related loss (El-Salhy et al., 1999; Wang et al., 2013). Age-related changes in Calretinin^+^ neurons have been described in the guinea pig and rat ileum (Abalo et al., 2005; Thrasivoulou et al., 2006) and in the mouse internal anal sphincter (Wang et al., 2013). We sought to extend this analysis of Calretinin^+^ neurons, a population that in the mouse myenteric plexus includes excitatory motor neurons (Drokhlyansky et al., 2020). We focused on the small intestine to assess whether age-related changes recapitulate in the ileum and thereby extend to adjacent intestinal regions. We represented Calretinin^+^ neurons as a percentage of HuC/D^+^ neurons, and found that the percentage of HuC/D^+^ neurons that were also Calretinin^+^ did not change with age in the duodenum, jejunum, or ileum (Figures 4A,B). Thus, the relative size of the Calretinin^+^ population is maintained with age throughout the small intestine.

**Figure 4.**
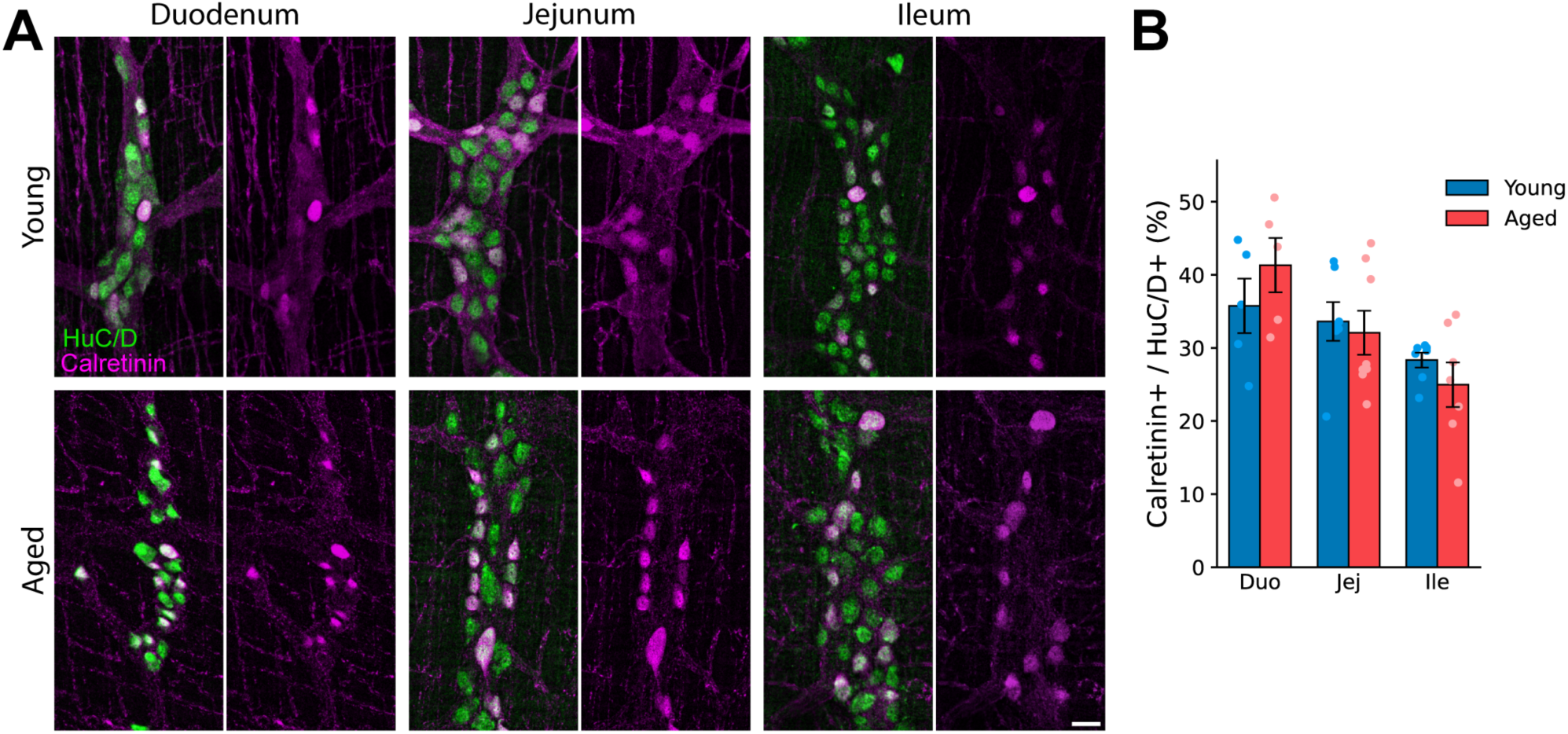
The proportion of Calretinin^+^ neurons does not change with age in the small intestine. **(A)** Representative HuC/D^+^ neuronal (green) and Calretinin^+^ (magenta) immunostaining in the duodenum, jejunum, and ileum of young (top) and aged (bottom) mice. **(B)** Percentage of HuC/D^+^ neurons that are Calretinin^+^ in the duodenum, jejunum, and ileum. Data are presented as mean ± SEM, with individual animals shown as points, *n* = 5-8 mice per group (8 young and 9 aged in total). Binomial GLMM followed by within-region comparisons of age. Scale bar, 25 µm. Duo, Duodenum; Jej, Jejunum; Ile, Ileum.

## 3 Discussion

Age-related changes in ENS cytoarchitecture have been reported in both animal models and humans, but with considerable variability between studies (Nguyen et al., 2023). Here, we compared myenteric plexus organization across all five regions of the small and large intestine in young and aged mice, and found that aging does not remodel the ENS uniformly. Neuronal density was unchanged with age in every region. While neurons in all regions maintain their stripe-like organization, the alignment of ganglia along these stripes is decreased in the duodenum and ileum. In the ileum and distal colon, ganglia were smaller and a greater proportion of neurons lay outside them. Glial number changed in a different set of regions, increasing in the duodenum and distal colon and decreasing in the proximal colon, while the proportion of glia within ganglia was maintained. Finally, the proportion of neurons that were Calretinin^+^ was unchanged across the small intestine. Together, these findings show that each region of the myenteric plexus follows its own regional pattern of change with age, and that in neurons, aging alters arrangement rather than density.

Most studies of the aging ENS have focused on neuron number and density, and most report a loss of myenteric neurons with age (Gomes et al., 1997; El-Salhy et al., 1999; Phillips et al., 2004; Abalo et al., 2005). However, Gamage et al. found that neuron numbers in the aged mouse distal colon were maintained once tissue stretch and an age-related increase in gut length and circumference were accounted for (Gamage et al., 2013). We found no evidence of neuron loss. Instead, ganglia in the ileum and distal colon contained fewer neurons, and more neurons were extraganglionic. This is consistent with earlier descriptions of reduced packing density in aged myenteric ganglia (El-Salhy et al., 1999; Abalo et al., 2005; Gamage et al., 2013). Because of our tissue preparation methodology, in which the GI tract was opened and maximally stretched before fixation, our measurements may not be directly comparable to studies of unstretched tissue, or to estimates of total neuron number. Importantly, however, the central conclusion of this study rests on comparisons between regions rather than absolute neuron counts. Tissue was prepared and analyzed identically, and moreover, because neuronal density was unchanged in every region, differential stretching of aged tissue is unlikely to account for changes in ganglionic organization. Most previous studies focus on only one or two regions, and our observed regional differences may help explain why reports of age-related neuronal loss are inconsistent.

The effects of aging on enteric glia have received less attention. In aged rats, glial density has been reported to decrease in the duodenum and ileum, while the ratio of glia to neurons increases in the duodenum (Phillips et al., 2004). In the aged human myenteric plexus, S100^+^ glia decrease in the descending colon (Baidoo et al., 2023) and GFAP^+^ glia decrease in the transverse colon (Ahmed et al., 2025). Here, we found that glial number changes in a region-specific manner, increasing in the duodenum and distal colon and decreasing in the proximal colon. The duodenal increase in glial density, with neuronal density unchanged, is in line with the higher glia-to-neuron ratio reported in aged rats, although Phillips et al. reported a net decrease rather than an increase in glia (Phillips et al., 2004). This difference may reflect their use of a different glial marker (S100) and a difference in species analyzed. Within each region, we found that the proportion of glia located within ganglia did not change, indicating that intraganglionic and extraganglionic glia were proportionately affected.

Constipation becomes more common with age in humans, and animal studies have shown reduced gastric and colonic motility with age (Patel et al., 2014; Saffrey, 2014). This makes the neurons that drive muscle contractions an important population to examine. Loss of cholinergic neurons, which include excitatory motor neurons, has been reported in the stomach, jejunum, and colon of aged mice and in the aged human colon (Bernard et al., 2009; Sun et al., 2018). Calretinin^+^ neurons, another population that includes excitatory motor neurons, have been examined in the ileum, where their numbers were unchanged with age in the rat and decreased in the guinea pig (Thrasivoulou et al., 2006; Abalo et al., 2007). We extended this analysis to all three regions of the mouse small intestine and found that the proportion of Calretinin^+^ neurons was maintained throughout. Because neuronal density was also unchanged, this suggests that Calretinin^+^ neurons are not lost with age in the small intestine. Whether this holds true in the colon, the region most directly linked to constipation, remains to be determined.

We acknowledge this study has several limitations. First, only male mice were used. Myenteric organization does not appear to differ between the sexes in young mice (Hamnett et al., 2022). However, constipation is more common in women (McCrea et al., 2009), and whether the ENS ages differently in females remains an open question. Second, we compared only two ages, so we cannot determine when the observed changes begin or how they progress. Finally, our analyses were structural. Whether the regional changes we describe affect motility in the corresponding regions of the GI tract will require further functional studies.

Aging affects the entire GI tract, from the oral cavity to the large intestine (Dumic et al., 2019). Although many studies have examined the aging ENS, most have focused on one or two regions, and differences in age, species, and quantification methods make their results difficult to compare. By comparing multiple features of myenteric organization across all regions of the small and large intestine, this study shows that the ENS does not age uniformly. Each region, and each component of the myenteric plexus, follows its own course. This regional map provides a framework for asking why some regions are more vulnerable to aging than others, and how region-specific changes in ENS organization relate to age-associated regional GI dysfunction.

## 4 Materials and Methods

### 4.1 Animals

All procedures were approved by the Stanford University Administrative Panel on Laboratory Animal Care and conformed to the National Institutes of Health *Guidelines for the Care and Use of Laboratory Animals*. Aged male C57BL/6 mice were obtained from the National Institute on Aging aged rodent colony maintained at Charles River Laboratories and were 22-26 months old at tissue collection. Young adult male C57BL/6 mice were obtained from Charles River Laboratories and were 3-4 months old at tissue collection. All mice were wild type. Mice were acclimated to the Stanford facility for at least 14 days before use. Mice were group housed (≤5 mice per cage) in individually ventilated cages under a 12h:12h light/dark cycle at 20-26°C and 30-70% relative humidity, with ad libitum access to water and food (Teklad Global 18% Protein Rodent Diet). Aged mice were inspected for general appearance and activity level before inclusion into experiments. A total of 8 young and 12 aged mice were used; sample sizes for each analysis are given in the figure legends.

### 4.2 Tissue Collection and Preparation

Intestinal dissections and tissue preparation were performed as previously described (Hamnett et al., 2022; Gomez-Frittelli et al., 2024). Mice were euthanized by CO_2_ inhalation followed by cervical dislocation. The intestines were removed and flushed of luminal contents with cold phosphate-buffered saline (PBS). A 2-3 cm segment was taken from the proximal end, the midpoint, and the distal end of the small intestine to obtain the duodenum, jejunum, and ileum, respectively. The colon was divided into proximal and distal halves. Segments were pinned to Sylgard 170-coated dishes in ice-cold PBS, the mesentery was trimmed away, and each segment was opened longitudinally along the mesenteric border. Tissue was then maximally stretched flat and pinned serosal side up and fixed in 4% paraformaldehyde for 90 min at 4°C with gentle shaking. After fixation, the muscularis externa, containing the myenteric plexus, was peeled from the underlying submucosa and mucosa with fine forceps and retained for whole-mount immunohistochemistry. Tissue was stored for no longer than 2 weeks at 4°C in PBS.

### 4.3 Immunohistochemistry

Immunohistochemistry was performed as previously described (Hamnett et al., 2022). Whole-mount muscularis preparations were cut into pieces of approximately 1 cm^2^ and placed in WHO microtitration trays (International Scientific Supplies) containing PBS. Tissue was blocked in a solution of PBS containing 5% normal donkey serum, 1% bovine serum albumin (BSA) and 0.1% Triton X-100 for 1 hour at room temperature with shaking. Tissue was then incubated with primary antibodies (Supplementary Table S1) diluted in PBT (PBS containing 1% BSA and 0.1% Triton X-100) overnight at 4°C with shaking. The following day, tissue was washed three times for 30 minutes each in PBT and then incubated with fluorophore-conjugated secondary antibodies (Supplementary Table S1) diluted in PBT for 2 hours at room temperature with shaking. After two 30 minute washes in PBT and two 30 minute washes in PBS, tissue was mounted onto glass slides using a fine paintbrush. Mounted tissue was air-dried, rinsed briefly in double-distilled water, and coverslipped in Fluoromount-G (SouthernBiotech).

### 4.4 Image Quantification

#### 4.4.1 Image Acquisition

Immunostained tissue was imaged on a Leica SP8 confocal microscope (Leica Microsystems) using a 20× oil objective (NA 0.75). Images were acquired at 1024 × 1024 pixels per tile (0.54 µm/pixel) with a 2 µm z-step, and z-stacks spanned the full depth of the myenteric plexus. For each gut region of each mouse, a single tile scan of approximately 3096 × 3022 µm was acquired for each staining experiment, avoiding any folded or damaged areas. Tile scans were collected with the Navigator module and stitched in LAS X (v1.4.7.28982), and maximum intensity projections were generated for analysis unless otherwise noted.

#### 4.4.2 Neuron and Ganglia Quantification

Single channel HuC/D images were exported to Fiji/ImageJ, maximum intensity projected and cropped to 1800 × 1800 µm fields. Neurons and ganglia were quantified with a modified version of COUNTEN (Kobayashi et al., 2021). We replaced the global threshold with local adaptive thresholding and background subtraction to accommodate variable staining intensity; full code and parameters available at https://doi.org/10.5281/zenodo.24120217. Ganglia were defined as clusters of ≥3 neurons (DBSCAN, ε = 25 µm), and unclustered neurons were classified as extraganglionic. Two anti-HuC/D antibodies were used (Supplementary Table S1, a rabbit monoclonal antibody and human anti-Hu serum), and both antibodies were represented in the young and aged groups. Because the human serum produced higher background, stricter thresholding parameters were used for human HuC/D-stained images. All other parameters were identical between antibodies. For each mouse and gut region, one or two fields were analyzed, and values were averaged where two were available. Segmentation outputs were visually inspected blind to age, and false-positive detections were removed.

#### 4.4.3 Stripe Analysis

Stripe organization was analyzed in 1800 × 1800 µm fields using the neuron positions and ganglia identified by COUNTEN (142 fields; 8 young and 12 aged mice). Stripes were traced by hand in Fiji as segmented lines following bands of ganglia. Before tracing, fields were randomized and renamed so that the tracer was blind to the age, mouse, and gut region of each field. Only segments that could be followed with confidence were traced (3-14 per field), and their coordinates were exported with a custom Fiji macro. Fields were not rotated, and all angles were measured in each field’s own image frame. The stripe direction of each field (φ) was the length-weighted mean direction of its traced segments, averaged as double angles because a line has no head or tail. The orientation of each ganglion (θ) was the long axis of an ellipse fitted to its segmented outline (scikit-image regionprops, van der Walt et al., 2014). Only ganglia of ≥8 neurons were analyzed (5,319 of 13,120 ganglia), because the long axis of smaller ganglia is poorly defined. The alignment of each ganglion with the stripes was quantified as cos 2(θ - φ), which equals 1 for a ganglion parallel to the stripe direction, 0 at 45°, and –1 for ganglia perpendicular to it. Each field was summarized by the mean alignment of its ganglia weighted by neuron count, and fields were averaged within each mouse and gut region. The decrease in alignment in the aged ileum was robust, persisting under a number of different conditions. These conditions included: changing the minimum ganglion size to 3, 5, or 12 neurons, weighting ganglia equally, comparing each ganglion with the direction of the nearest traced segment instead of the field-wide direction, and analyzing ganglia of 8–11, 12–19, and ≥20 neurons separately. Code is available at https://doi.org/10.5281/zenodo.24120217.

#### 4.4.4 Sox10 Analysis

Sox10 images were analyzed using Fiji/ImageJ. Maximum intensity projections were cropped to 1800 × 1800 µm fields and contrast was normalized (Enhance Contrast, 1% saturated pixels). Background was removed by a rolling ball subtraction (radius 50 px) and images were thresholded with a MaxEntropy algorithm. The resulting binary masks were processed with hole filling, a single erosion and dilation, and watershed separation of touching nuclei. Sox10^+^ nuclei were then counted with Analyze Particles, excluding objects smaller than 40 px (≈12 µm^2^). To count intraganglionic glia, COUNTEN was re-run on the HuC/D channel of each Sox10 field to generate a matched ganglion mask (see section 4.4.2), which was combined with the Sox10 mask using a logical AND operation. The Sox10^+^ nuclei within ganglia were then counted. Extraganglionic glia were calculated as the total minus the intraganglionic count.

#### 4.4.5 Calretinin Analysis

Calretinin^+^ and HuC/D^+^ neurons were counted in the same 1800 × 1800 µm fields with a custom Python pipeline, available at https://doi.org/10.5281/zenodo.24120217. Calretinin^+^ cells were segmented by Gaussian smoothing, contrast normalization, and rolling-ball background subtraction, followed by local adaptive thresholding (400 µm neighborhood) with a floor anchored to the image’s MaxEntropy threshold. Touching cells were separated by watershed, and objects smaller than 58.5 µm^2^ or touching the image border were excluded. HuC/D^+^ neurons were detected with the approach described in section 4.4.2, using parameters re-optimized for these images. All parameters were defined in micrometers and converted to pixels for each image using its recorded pixel size.

Because Calretinin staining varied in intensity between samples, the threshold floor and overall threshold were adjusted per image with an interactive tool. Images were presented in randomized order, blind to age. Regions of a field that were unusable because of tissue tears or folds, dim staining, or bright non-somatic fiber signal were then excluded by selecting a rectangular sub-region, again in randomized order and blind to age, with counts hidden until the selection was committed. Calretinin^+^ and HuC/D^+^ neurons were counted within the retained region. Calretinin^+^ neurons were expressed as a percentage of HuC/D^+^ neurons in the same region, which controlled for differences in tissue area.

### 4.5 Statistical Analyses

Statistical analyses were performed in JMP Student Edition 19 (SAS Institute). The experimental unit was an individual mouse. For neuron and ganglion measures (Figures 1 and 2), values from the one or two fields analyzed per mouse and gut region were averaged; all other analyses used one field per mouse and region. All models included age, gut region, and their interaction as fixed effects, with mouse as a random effect to account for repeated measurements across regions within each mouse. The model family was chosen to suit each outcome. Continuous outcomes were analyzed with linear mixed models: numbers of neurons, ganglia, and extraganglionic neurons per field, as well as ganglion–stripe alignment, weighted by the number of scored ganglia. Mean and 90^th^-percentile neurons per ganglion and mean ganglion area (weighted by number of ganglia) were log_10_-transformed to reduce heteroscedasticity. Counts (total, intraganglionic, and extraganglionic Sox10^+^ cells) were analyzed with Poisson generalized linear mixed models (GLMMs). Proportions (extraganglionic neurons among all neurons, intraganglionic Sox10^+^ cells among all Sox10^+^ cells, Calretinin^+^ neurons among HuC/D^+^ neurons) were analyzed with binomial GLMMs with a logit link, using the denominator count as the number of trials. All Poisson and binomial GLMMs included an additional mouse × region random effect to account for overdispersion. Young and aged mice were compared within each gut region by slice tests of the age × region interaction; these within-region comparisons were not corrected across regions.

To analyze the distribution of ganglion sizes, ganglia from all fields of each mouse and region were pooled and binned by neurons per ganglion and by ganglion area, with bin edges set at the tertiles of the distribution pooled across all animals and regions. For each bin, the number of ganglia in the bin out of the total number of ganglia was analyzed with a binomial GLMM (logit link) with the same fixed effects and with mouse and mouse × region as random effects. Age was tested within each region by slice tests, and p-values were corrected with the Benjamini-Hochberg procedure separately for each metric (15 tests each: 5 regions × 3 bins; q<0.05).

Data are presented as mean ± SEM, with individual animals as points. Statistical significance was set at α = 0.05. For all analyses, *p<0.05, **p<0.01, ***p<0.001. Sample sizes are given in the figure legends. Full test results are provided in Supplementary Table S2.

## Statements

### Author Contributions

JHM: Conceptualization, Data curation, Formal analysis, Investigation, Methodology, Software, Validation, Visualization, Writing – original draft, Writing – review & editing. SD: Data curation, Formal analysis, Methodology, Software, Writing – review & editing. LSB: Conceptualization, Resources, Writing – review & editing. JAK: Conceptualization, Data curation, Funding acquisition, Supervision, Validation, Writing – original draft, Writing – review & editing.

## Acknowledgements

We would like to thank all members of the Kaltschmidt lab for their valuable feedback. We thank V. Lennon (Mayo Clinic) for the HuC/D antibody. We acknowledge the use of generative AI technology (Claude, Opus 5.5; Anthropic) to proofread the manuscript text, check for adherence to journal guidelines, and generate and edit image-analysis code. All AI-assisted content was reviewed and verified by the authors, including for factual accuracy and ensuring plagiarism-free content.

## Data Availability

The raw data supporting the conclusions of this article will be made available by the authors, without undue reservation. The code for neuron and ganglia detection, stripe analysis, and Calretinin quantification is available at Zenodo (https://doi.org/10.5281/zenodo.24120217)

## Ethics Statement

The animal study was approved by Stanford University Administrative Panel on Laboratory Animal Care. The study was conducted in accordance with the local legislation and institutional requirements.

---

**Figure S1.**
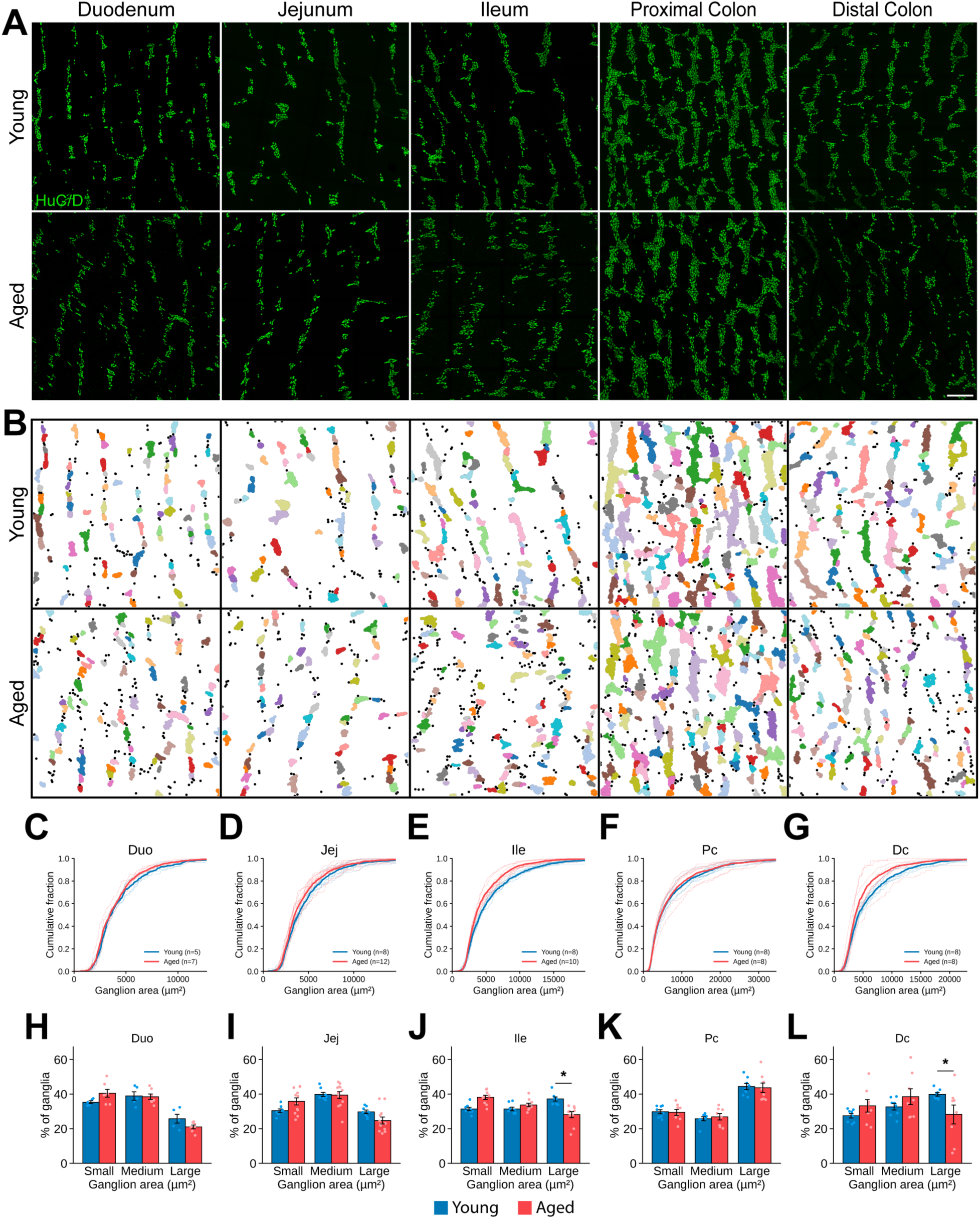
Ganglion size decreases with age in a region-specific manner. **(A)** Representative HuC/D^+^ neuronal immunostaining (green) across the length of the small and large intestines in young (top) and aged (bottom) mice. **(B)** Ganglia masks of the images in (A). Ganglia (clusters of ≥3 neurons, see Methods) are shown in distinct colors and extraganglionic neurons as black dots. **(C-G)** Empirical cumulative distribution functions (ECDFs) of ganglion area in young and aged mice in the duodenum **(C)**, jejunum **(D)**, ileum **(E)**, proximal colon **(F)**, and distal colon **(G)**, shown descriptively; statistical comparisons are presented in (H-L). **(H-L)** Percentage of ganglia classified as small (<2962 µm^2^), medium (2962-5311 µm^2^), or large (≥5311 µm^2^) in the duodenum **(H)**, jejunum **(I)**, ileum **(J)**, proximal colon **(K)**, and distal colon **(L)**. Size classes were defined by the tertiles of ganglion area pooled across all animals and regions. Data are presented as mean ± SEM, with individual animals shown as points; *n* = 5-12 mice per group (8 young and 12 aged in total). (H-L) Binomial GLMMs followed by within-region comparisons of age with Benjamini-Hochberg correction. *q<0.05. Scale bar, 250 µm. Duo, Duodenum; Jej, Jejunum; Ile, Ileum; Pc, Proximal colon; Dc, Distal colon.

**Figure S2.**
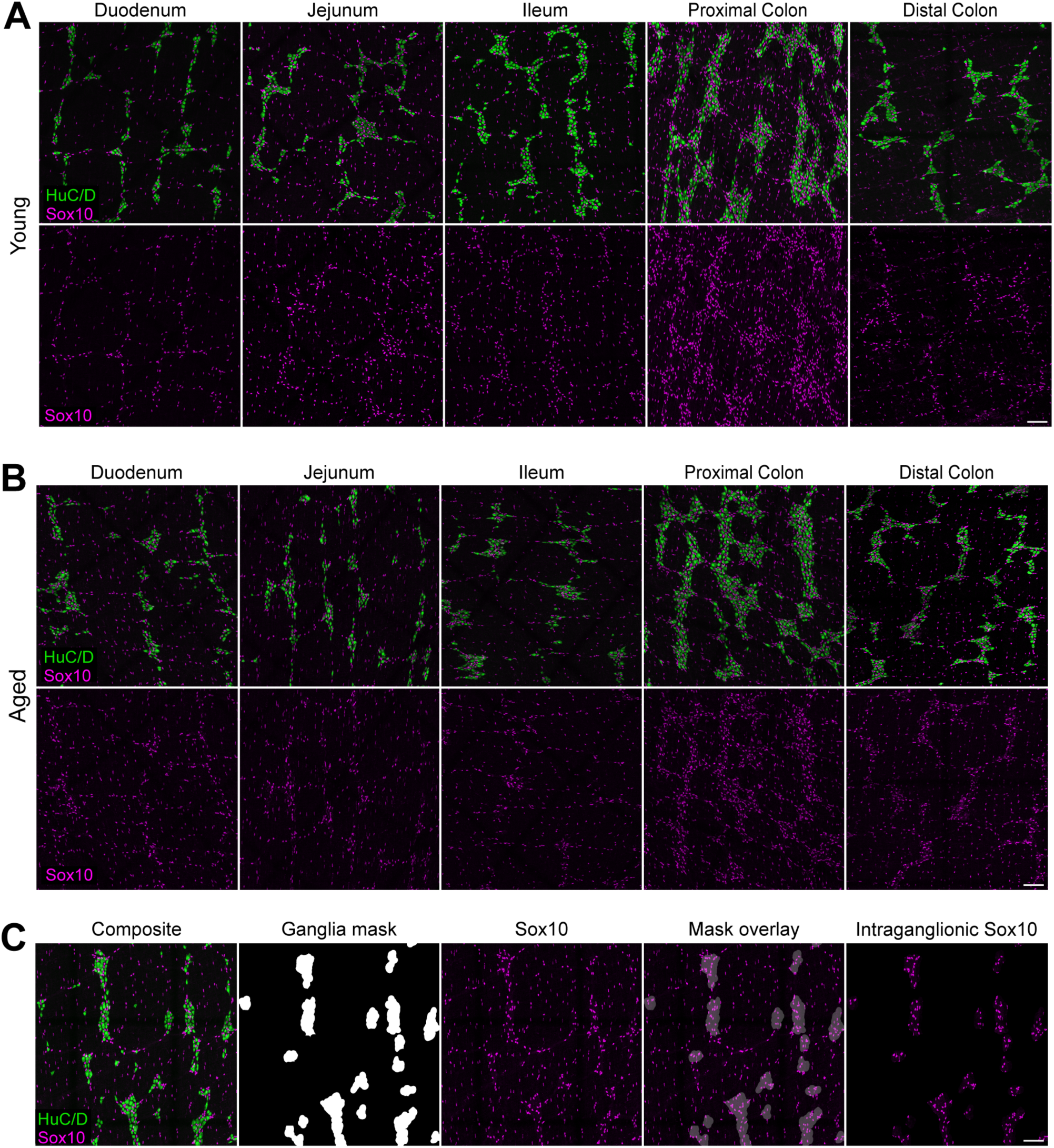
Glial distribution across the gut. **(A, B)** Representative HuC/D^+^ neuronal (green) and Sox10^+^ glial (magenta) immunostaining across all regions of the small and large intestine in young (A) and aged (B) mice. **(C)** Pipeline for identifying intraganglionic Sox10^+^cells. From left to right: raw composite image; ganglia masks (see Figure 2 and Methods); Sox10^+^ cells; ganglia mask overlaid on Sox10^+^ cells; and intraganglionic Sox10^+^ cells overlapping a ganglia mask. Scale bars, 100 µm.

**Supplementary Table S1.** Primary and secondary antibodies used.

| **Antigen** | **Host** | **Source** | **Catalog #** | **RRID** | **Dilution** |
|---|---|---|---|---|---|
| HuC/D (ANNA-1 serum) | Human | Gift of V. Lennon (Mayo Clinic) | N/A | N/A | 1:100,000 |
| HuC/D | Rabbit | Abcam | ab184267 | AB_2864321 | 1:4,000 |
| Sox10 | Goat | R&D Systems | AF2864 | AB_442208 | 1:2,000 |
| Calretinin | Goat | Swant | CG1 | AB_10000342 | 1:8,000 |
| Anti-Rabbit AF 488 | Donkey | Invitrogen | A21206 | AB_2535792 | 1:800 |
| Anti-Rabbit Cy3 | Donkey | Jackson Immuno Research | 711-165-152 | AB_2307443 | 1:800 |
| Anti-Human AF 488 | Donkey | Jackson Immuno Research | 709-545-149 | AB_2340566 | 1:800 |
| Anti-Goat Cy3 | Donkey | Jackson Immuno Research | 705-165-147 | AB_2307351 | 1:800 |
| Anti-Goat Cy5 | Donkey | Jackson Immuno Research | 705-175-147 | AB_2340415 | 1:800 |

**Supplementary Table S2.** Statistical tests for all quantified panels. Young and aged mice were compared within each gut region.

| **Figure** | **Measure** | **Statistical test** | **Transformation / weighting / offset** | **Duodenum** |  | **Jejunum** |  | **Ileum** |  | **Proximal colon** |  | **Distal colon** |  |
|---|---|---|---|---|---|---|---|---|---|---|---|---|---|
|  |  |  |  | **\*** | ***p*** | **\*** | ***p*** | **\*** | ***p*** | **\*** | ***p*** | **\*** | ***p*** |
| 1C | Neurons per field | LMM | None | ns | .9129 | ns | .8521 | ns | .9306 | ns | .9213 | ns | .4597 |
| 1D | Ganglion–stripe alignment<sup>c</sup> | LMM | Weighted by number of scored ganglia | * | .0401 | ns | .0842 | *** | <.0001 | ns | .1689 | ns | .3490 |
| 2C | Ganglia per field | LMM | None | ns | .6146 | ns | .9783 | ns | .0553 | ns | .6293 | ns | .7335 |
| 2D | Extraganglionic neurons per field | LMM | None | ns | .3517 | ns | .0742 | *** | .0009 | ns | .4319 | *** | .0007 |
| 2E | Extraganglionic neurons (% of neurons) | Binomial GLMM | Trials = total neurons | ns | .4189 | ns | .0606 | ** | .0060 | ns | .5454 | *** | .0002 |
| 2F | Mean ganglion area | LMM | log<sub>10</sub>; weighted by number of ganglia | ns | .4372 | ns | .1801 | ** | .0035 | ns | .2333 | ** | .0041 |
| 2G | Mean neurons per ganglion | LMM | log<sub>10</sub> | ns | .6085 | ns | .2585 | ** | .0052 | ns | .5169 | ** | .0011 |
| 2H | 90<sup>th</sup>-percentile neurons per ganglion | LMM | log<sub>10</sub> | ns | .3889 | ns | .4282 | ** | .0043 | ns | .3690 | *** | .0006 |
| 2I,J<sup>a</sup> | Ganglia with 3–4 neurons (% of ganglia) | Binomial GLMM | Trials = total ganglia; *q*<sup>b</sup> | ns | .8449 | ns | .1515 | ns | .1054 | ns | .8449 | * | .0120 |
| 2I,J<sup>a</sup> | Ganglia with 5–9 neurons (% of ganglia) | Binomial GLMM | Trials = total ganglia; *q*<sup>b</sup> | ns | .3516 | ns | .7288 | ns | .1116 | ns | .6686 | ns | .6686 |
| 2I,J<sup>a</sup> | Ganglia with ≥10 neurons (% of ganglia) | Binomial GLMM | Trials = total ganglia; *q*<sup>b</sup> | ns | .3562 | ns | .1860 | * | .0120 | ns | .7288 | * | .0120 |
| S1H–L | Small, <2962 µm² (% of ganglia) | Binomial GLMM | Trials = total ganglia; *q*<sup>b</sup> | ns | .3502 | ns | .1170 | ns | .0595 | ns | .9488 | ns | .1236 |
| S1H–L | Medium, 2962–5311 µm² (% of ganglia) | Binomial GLMM | Trials = total ganglia; *q*<sup>b</sup> | ns | .9488 | ns | .9171 | ns | .3675 | ns | .8951 | ns | .3502 |
| S1H–L | Large, ≥5311 µm² (% of ganglia) | Binomial GLMM | Trials = total ganglia; *q*<sup>b</sup> | ns | .3502 | ns | .1575 | * | .0217 | ns | .9171 | * | .0217 |
| 3C | Sox10<sup>+</sup> cells per field | Poisson GLMM | None | * | .0236 | ns | .2033 | ns | .3911 | * | .0349 | ** | .0096 |
| 3D | Intraganglionic Sox10<sup>+</sup> cells per field | Poisson GLMM | None | ns | .4011 | ns | .2321 | ns | .4902 | ns | .2407 | * | .0444 |
| 3E | Extraganglionic Sox10<sup>+</sup> cells per field | Poisson GLMM | None | ** | .0098 | ns | .3131 | ns | .5575 | ** | .0069 | * | .0403 |
| 3F | Intraganglionic Sox10<sup>+</sup> cells (% of Sox10<sup>+</sup>) | Binomial GLMM | Trials = total Sox10<sup>+</sup> cells | ns | .2652 | ns | .6503 | ns | .7686 | ns | .3628 | ns | .6748 |
| 4B | Calretinin<sup>+</sup> neurons (% of HuC/D<sup>+</sup>) | Binomial GLMM | Trials = total HuC/D<sup>+</sup> neurons | ns | .1778 | ns | .8289 | ns | .2587 | — |  | — |  |

All models included age, gut region, and age × region as fixed effects and mouse as a random effect. Poisson and binomial GLMMs also included a mouse × region random effect to account for overdispersion. Young and aged mice were compared within each region by slice tests of the age × region interaction; within-region comparisons were not corrected across regions. Neuron and ganglion measures (Figures 1, 2, and S1) were averaged over the one or two fields per mouse and region (ganglion-size bins pooled ganglia across fields); all other analyses used one field per mouse and region.
<sup>a</sup> Figure 2I,J show the duodenum and ileum; all five regions were tested. <sup>b</sup> *q* values: Benjamini–Hochberg false discovery rate correction across 15 tests per metric (5 regions × 3 bins); the *p* column reports *q* values for these rows. <sup>c</sup> Alignment of the long axis of each ganglion of ≥8 neurons with the traced stripe direction of its field, from −1 (perpendicular) to +1 (parallel); see Methods.
—, not analyzed (colon excluded from the calretinin analysis). ns, not significant; \**p* < 0.05; \*\**p* < 0.01; \*\*\**p* < 0.001. LMM, linear mixed model; GLMM, generalized linear mixed model.

